# Predicting aging of brain metabolic topography using variational autoencoder

**DOI:** 10.1101/200865

**Authors:** Hongyoon Choi, Hyejin Kang, Dong Soo Lee, Alzheimer’s Disease Neuroimaging Initiative

## Abstract

Predicting future brain topography can give insight into neural correlates of aging and neurodegeneration. Due to variability in aging process, it has been challenging to precisely estimate brain topographical change according to aging. Here, we predict age-related brain metabolic change by generating future brain ^18^F-Fluorodeoxyglucose PET. A cross-sectional PET dataset of cognitively normal subjects with different age was used to develop a generative model. The model generated PET images using age information and characteristic individual features. Predicted regional metabolic changes were correlated with the real changes obtained by follow-up data. This model was applied to produce a brain metabolism aging movie by generating PET at different ages. Normal population distribution of brain metabolic topography at each age was estimated as well. In addition, a generative model using APOE4 status as well as age as inputs revealed a significant effect of APOE4 status on age-related metabolic changes particularly in the calcarine, lingual cortex, hippocampus and amygdala. It suggested APOE4 could be a factor affecting individual variability in age-related metabolic degeneration in normal elderly. This predictive model may not only be extended to understanding cognitive aging process, but apply to development of a preclinical biomarker for various brain disorders.

## Introduction

Understanding the normal aging change in the brain is essential to understand neural correlates of cognitive aging and to investigate various neurodegenerative diseases including Alzheimer’s disease (*1*). In particular, brain metabolism which can be measured by ^18^F-fluorodeoxyglucose (FDG) PET has been regarded as a key biomarker for neurodegenerative disorders. Identifying brain metabolic topography associated with aging could give insight into the neural basis of age-related cognitive decline and help differentiate normal aging from neurodegenerative disorders.

Although the relationship between cerebral glucose metabolism and aging has been repeatedly studied, there has been controversy about which brain regions show significant age-related metabolic decline (*2–6*). Individual genetic background and healthy status as well as underlying brain disease gives rise to the individual variability in age-related metabolic change (*7*, *8*). Due to this variability, we have not been able to predict individual aged brain understandably. Therefore, instead of consideration of individual variability, previous studies have focused on the trend of overall aging changes using cross-sectional imaging data with statistical models such as linear regression. Even though this statistical analysis could provide overall brain metabolic changes, it was difficult to individually apply to estimate how far a given subject’s brain metabolism is from the normal population at the same age. This individual evaluation of brain metabolism can be extended to the differentiation between normal and abnormal aging process. It requires normal population distribution database of all ages, however, it has been challenging to build a database of the population distribution of normal brain metabolism for each age from the cross-sectional data with subjects of various age distribution.

Here, we develop a model for predicting future brain metabolic topography by generating brain PET image. In this study, we utilize variational autoencoder (VAE), a type of unsupervised learning methods, which can generate images from some representations (VAE) (*9*). We applied it to predicting FDG brain PET at different ages. Each FDG PET image combined with the subject’s current age information was represented by low-dimensional features and then PET images corresponding different ages were generated. We also generated population distribution data of normal brain metabolic topography at different ages, which represented variability om individual metabolic activity at each age. As an application of our approach to discovering factors that potentially affect brain aging, we further investigated whether APOE4 status impacted on the age-related metabolic change by using a generative model that uses age and APOE4 information.

## Results

### Prediction of future brain metabolic change

The VAE-based model was designed to represent FDG PET images and corresponding subjects’ age to latent features (**Fig. 1A**). The posterior part of this model, a generator component, could produce PET images from any values of the latent features and age information. The model was trained by baseline PET images of 393 cognitively normal subjects.

**Figure 1.**
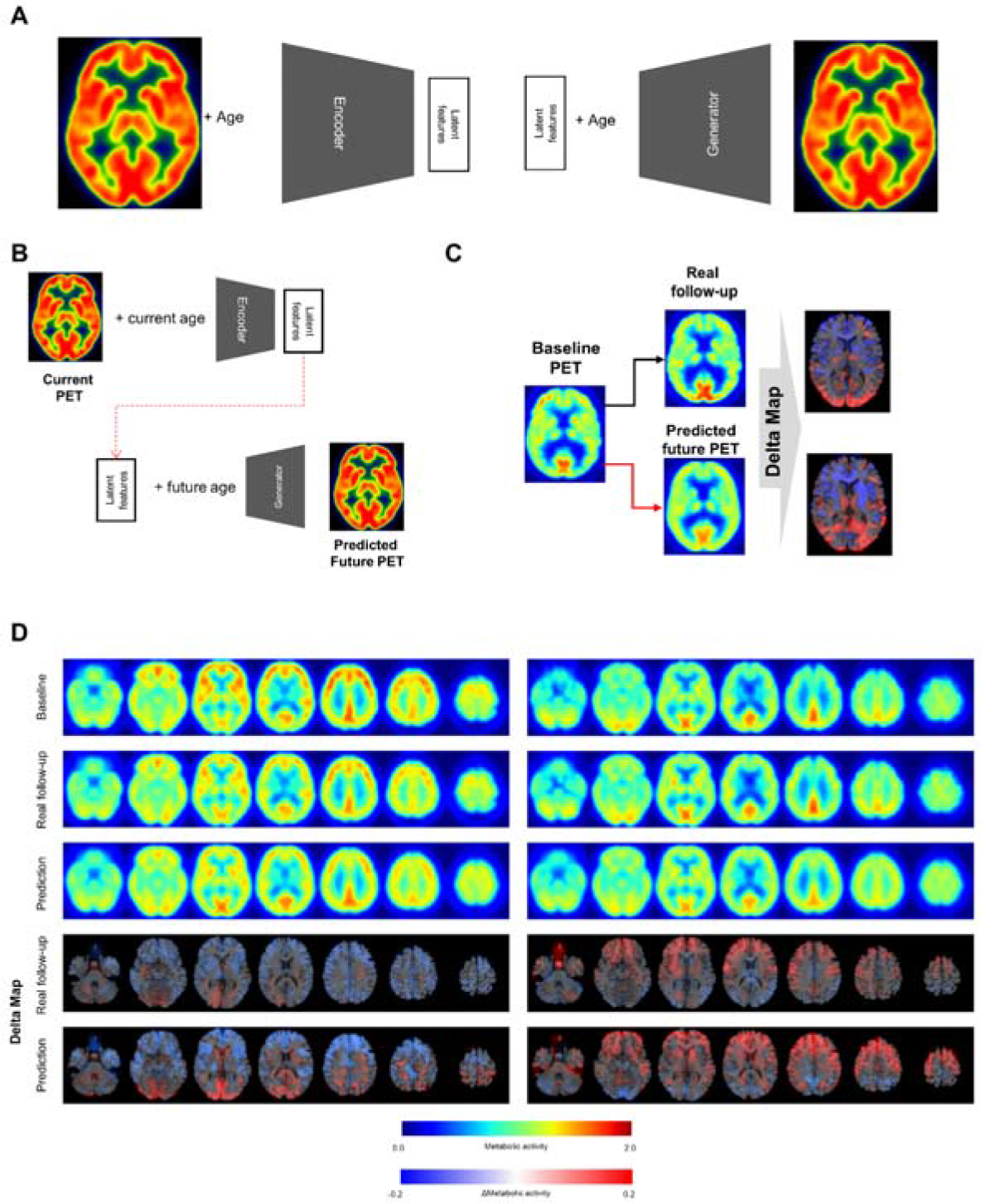
Metabolic change prediction by generating future brain PET. (A) VAE model which consists of encoder and generator was trained by PET images of cognitively normal subjects. The encoder represents input PET images to 10 latent features. The generator generates virtual PET image from any values of latent features and age information. (B) The VAE-based model could generate future brain PET individually using baseline PET image. A subject’s brain PET was encoded into latent features. We hypothesized that these latent features were unchanged across age. Future brain PET was generated by entering future age and the latent features. (C) Predicted individually generated PET was compared with real follow-up data. For comparison, delta maps obtained by subtracting baseline from prediction or follow-up images were generated. (D) Representative cases follow-up PET and individually predicted PET. According to the follow-up data, there was comparable individual variability in metabolic change. A subject showed globally decreased metabolism (left) while another subject showed increased metabolism in the frontotemporal cortex (right). Predicted future PET could also reflect the individual variability.

To generate future brain PET images, we firstly obtained latent features of a subject’s baseline PET image using the encoder. We assumed that these were not changed according to aging as characteristic individual features. The features of a subject were entered into the generator with any age, which could generate the subject’s virtual brain PET at different ages (**Fig. 1B**). The model was tested by cognitively healthy subjects who underwent both baseline and follow-up PET. Follow-up PET scans were obtained for 26 subjects after 4 years from baseline and for 11 subjects after 5 years. Predicted metabolic change was compared with corresponding real metabolic change computed by follow-up PET data. Each predicted future brain PET and real follow-up PET was subtracted by corresponding baseline PET for the comparison (**Fig. 1C**). As a result, delta maps, the future brain PET subtracted by baseline, obtained from real follow-up PET showed individual variability. Corresponding predicted future brain PET also showed those variable patterns (**Fig. 1D**). A subject showed prominently decreased metabolism in the cerebral cortices, while another showed relatively increased metabolism in the frontal cortex (**Fig. 1D**). The delta map obtained by real follow-up was positively correlated with that obtained by prediction (**fig. S1**).

To compare predicted future brain PET and real follow-up PET quantitatively, mean metabolic changes of 116 predefined brain regions across all subjects were calculated. Averaged predicted changes in regional metabolism was significantly correlated with the real changes obtained by real follow-up data (r= 0.59, p < 0.001 and r=0.59, p < 0.001 for 4-year and 5-year follow-up, respectively; **Fig. 2A and B**). Bland-Altman plot showed the difference between predicted and real regional metabolic activities (**Fig. 2C and D**). The 95% confidence interval of the prediction error of regional metabolic activity was -0.027 – 0.027 for 4-year follow-up and -0.027 – 0.048 for 5-year follow-up. In addition, individually predicted and real metabolic changes were compared. To show how individual prediction of metabolic change was similar with the real change, voxelwise correlations of individual delta maps obtained by follow-up and prediction were calculated. We could find a trend of high correlation between the two delta maps of same subject though the prediction of metabolic change was failed in some subjects (**fig. S2**). The similarity between predicted and real metabolic change was not significantly affected by subjects’ age, gender, follow-up diagnosis, APOE4 and baseline mini-mental state examination.

**Figure 2.**
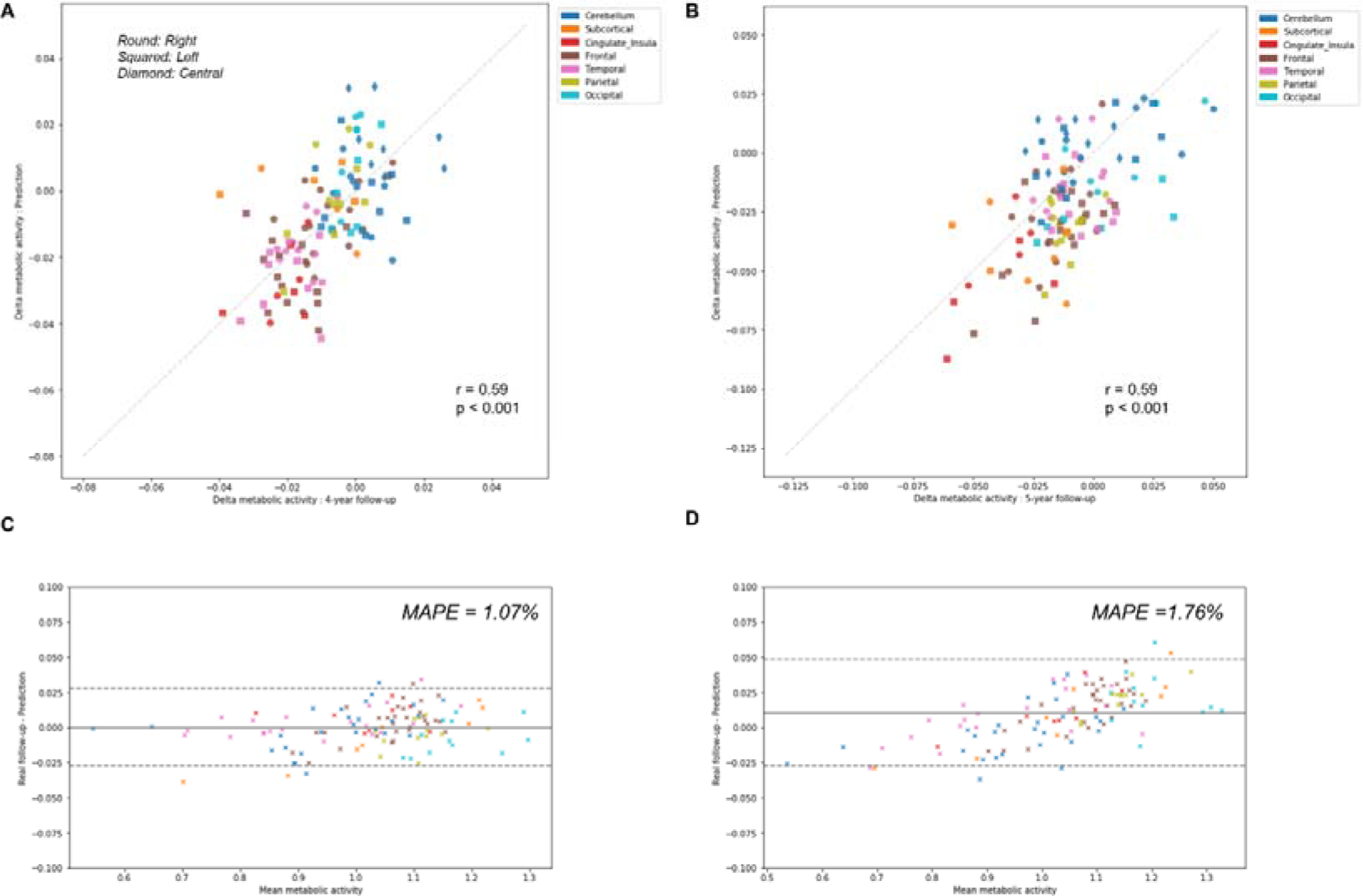
Comparison of predicted metabolic change with real follow-up data. Regional metabolic change from baseline was averaged across subjects for predicted and follow-up data. Averaged predicted and real changes across the brain regions were significantly correlated for 4-year follow-up images (r = 0.59, p < 0.001) (A) and 5-year follow-up images (r = 0.59, p < 0.001) (B). Bland-Altman plots were drawn for the comparison of predicted and real regional metabolic activity for 4-year (C) and 5-year PET images (D). The 95% confidence interval of the error of predicted regional metabolism was -0.027 – 0.027 for 4-year follow-up and -0.027 – 0.048 for 5-year follow-up. MAPE was 1.07% for 4-year follow-up and 1.76% for 5-year follow-up.

### Generating overall brain metabolism aging movie

We applied our model to the assessment of overall regional metabolic changes. To investigate overall patterns of age-related brain metabolism, representative brain images were generated by using different age and mean value of each latent feature across all subjects (**Fig. 3A**). The representative FDG brain PET generated from the age of 50 to 90 is presented in **Fig. 3B**. To visualize the age-related change definitely, the generated FDG PET with different age was subtracted by generated PET of the age of 50 (**Fig. 3C and D; fig. S3**).

**Figure 3.**
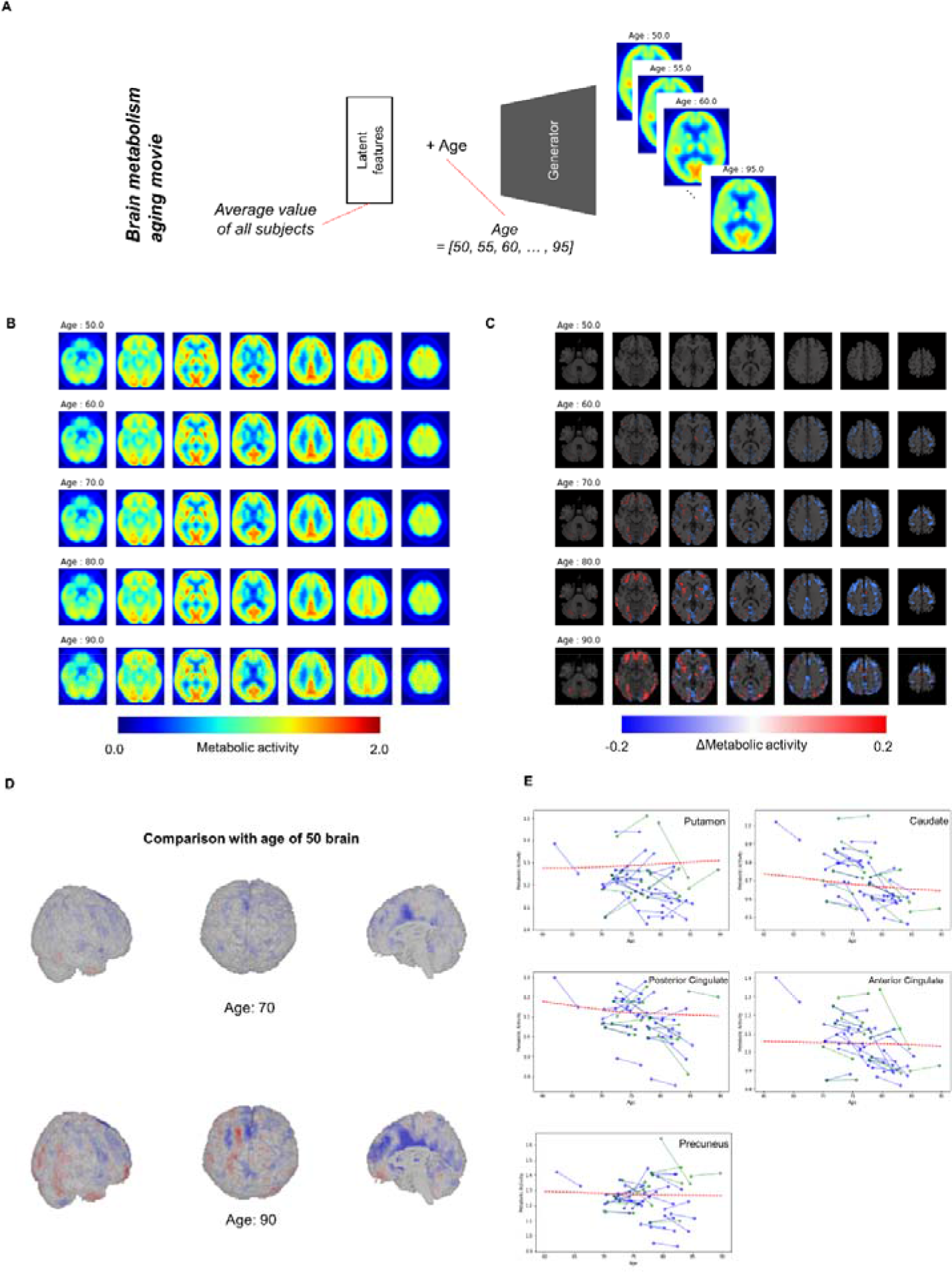
Overall brain metabolism aging movie by generating representative PET of each age. (A) Using VAE-based model, representative FDG PET images of different age were generated to identify overall age-related metabolic pattern. Mean latent feature values across all trained subjects were entered into generator for representative PET images. (B) Using mean latent features, representative PET images were generated according to aging. (C) Compared with the representative PET of age of 50, subtraction images were generated. (D) Surface visualization of the subtraction map revealed that age-related decline was mainly found in the cingulate cortex. (E) Age-related metabolic change in specific brain regions was plotted. Solid lines represent real metabolic change data for 4-year follow-up (blue) and 5-year follow-up (green). Red dotted lines represent regional metabolic changes estimated by virtually generated PET images.

**Fig. 3D** showed that age-related metabolism decline was mainly found in the cingulate cortex. Using predefined brain regions of interests, metabolic activity of each brain region was extracted according to aging (**Fig. 3E**). Red dotted lines represent estimated metabolic decline using the generated PET by entering mean latent features. Solid lines represent real metabolic decline obtained by 4-year (Blue) and 5-year (Green) follow-up data (**Fig. 3E**). The curves estimated by the VAE model explained that overall metabolic decline with aging was nonlinear. Approximately before 75, age-related metabolic decline was steep in the posterior cingulate and caudate and then the decline became slower after 75.

### Distribution of regional metabolic activity at each age

Most brain imaging data including our subjects consist of imaging with various ages. Thus, it has been hard to obtain population distribution of normal brain at each age. Randomly resampled latent features could generate population distribution of regional brain metabolic activity for all ages (**Fig. 4A**). Generated brain PET data from resampled latent features provide the variety of regional metabolic activity. Histograms of each brain region at different ages were drawn (**Fig. 4B**). As aforementioned representative brain metabolic changes, histograms of posterior cingulate and caudate showed a trend of left shifting according to aging. Distribution of overall aging patterns of regional metabolism was also exhibited (**Fig. 4C, fig. S4**). Dotted lines represent 95% confidence intervals of regional metabolic activity.

**Figure 4.**
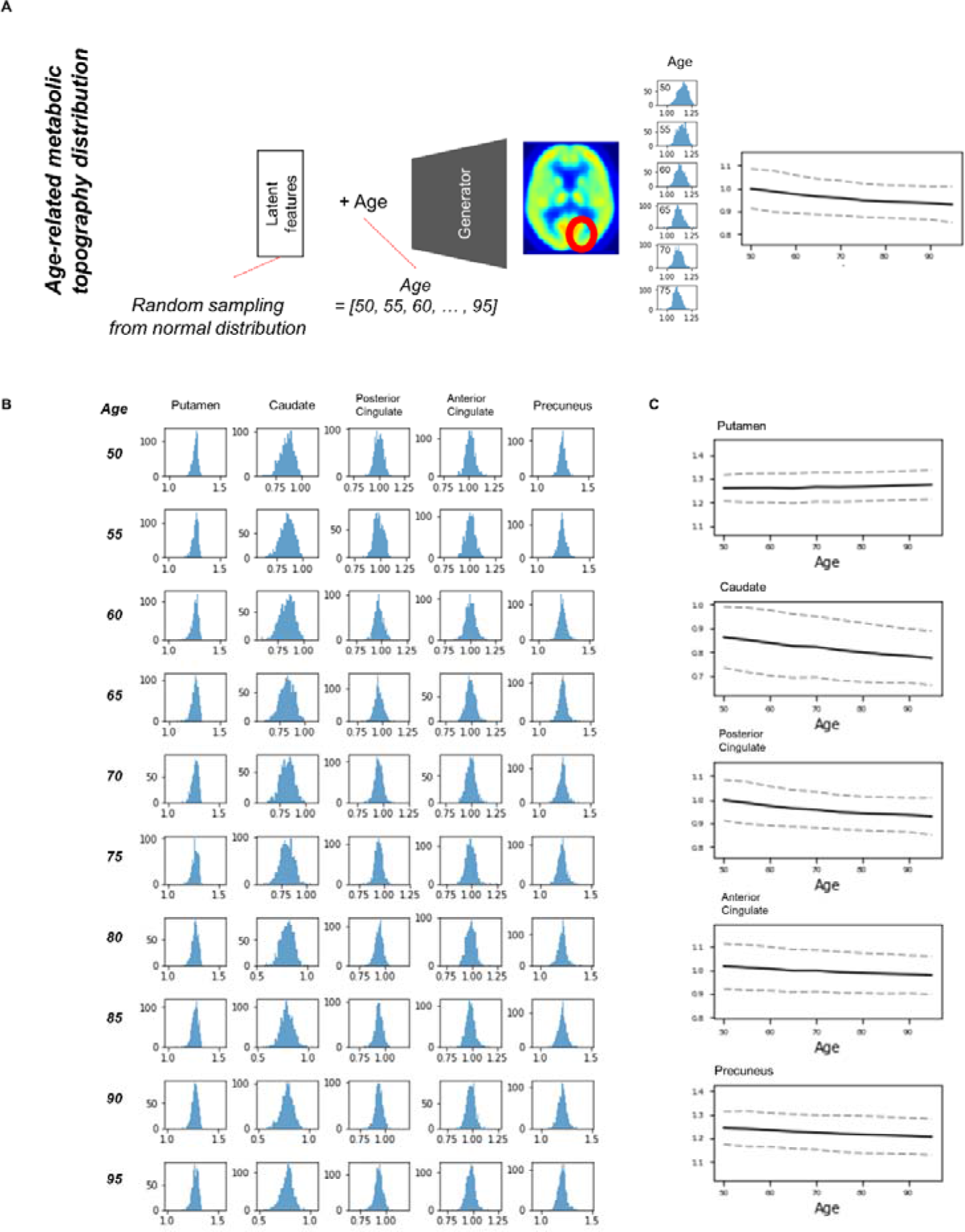
Estimating population distribution of brain metabolism at each age. (A) Population distribution of brain metabolic topography was estimated by resampling latent features. Generated brain PET was repeatedly generated by random latent feature values sampled from normal distribution. Distribution of regional metabolism was estimated for all ages. (B) Histograms of distribution of metabolic activity were drawn for putamen, caudate, posterior cingulate, anterior cingulate and precuneus at different ages. (C) Confidence intervals of metabolic changes could be estimated by the distribution. Dotted lines represent 95% confidence interval of regional metabolic activity.

We found individual variability in regional brain metabolism at different ages. The individual variability was determined by the distribution of latent features. To show how each latent feature affects brain metabolism, PET images were generated by changing latent features. Brain metabolic patterns were changed according to latent features as shown in **Fig. 5**. As an example, increased feature 1 was associated with decreased brain metabolism in the posterior temporal and occipital cortices and increased feature 2 was associated with increased frontal metabolism.

**Figure 5.**
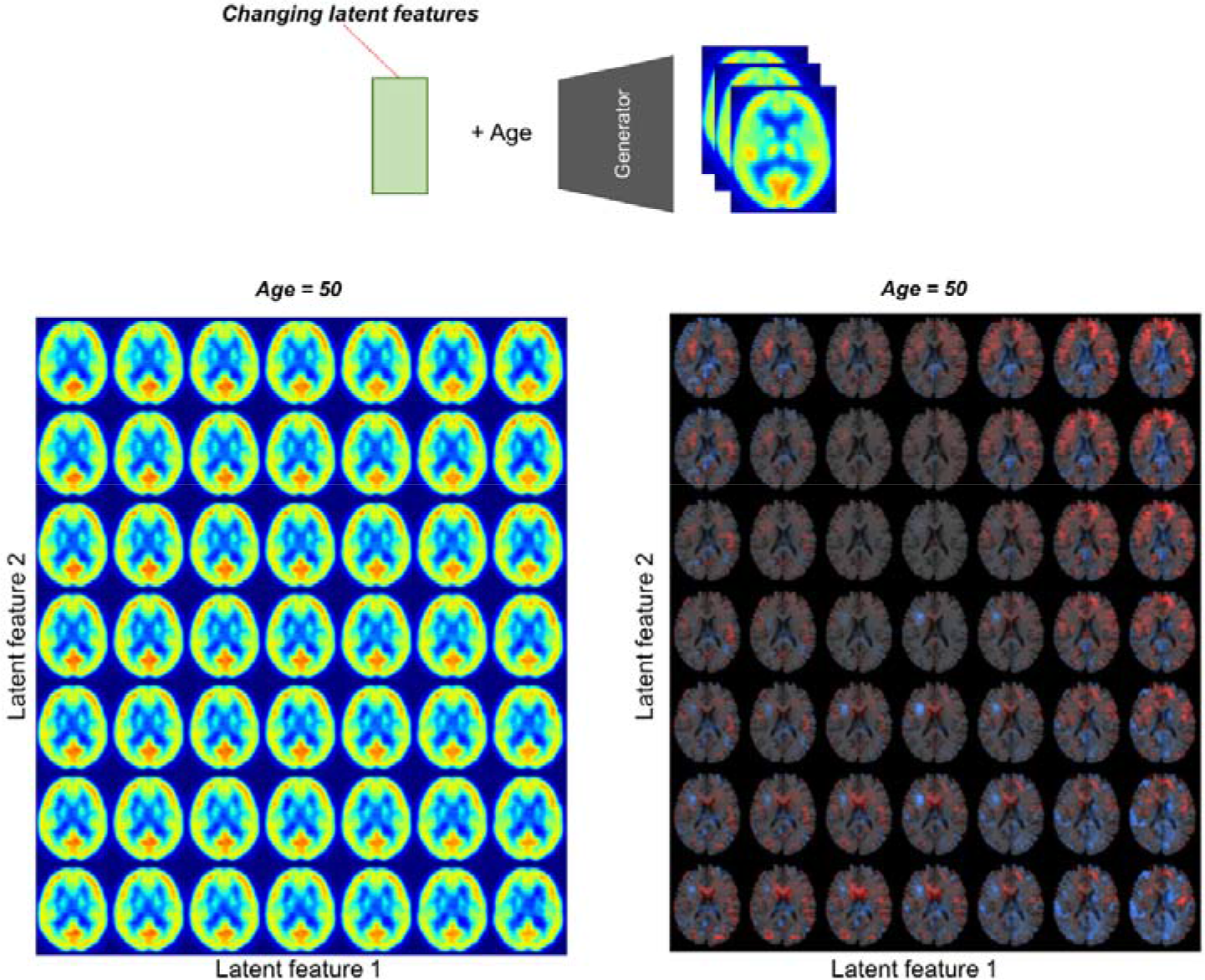
Brain metabolic topography according to latent features. As the encoder of VAE compressed PET image into 10 latent features, variability in brain metabolism is determined by these 10 features. To assess metabolic patterns determined by latent features, brain PET images were generated according to different latent feature values. An example of the two latent features, increased first latent feature (x-axis) was associated with decreased metabolism in posterior temporal and occipital cortices. Increased second latent feature (y-axis) was associated with increased metabolism in the frontotemporal cortices at age of 50.

### APOE4 status and age-related metabolic change

Because clinical variables affect age-related metabolic change and its variability, we further investigated whether APOE4 status impacts on metabolic changing patterns. Another VAE model was trained using two conditions, age and APOE4 status (**Fig. 6A**). This model can generate virtual brain PET images according to the age and APOE4 status. Thus, age-related metabolic change according to APOE4 can be estimated by inputting APOE4 positive and negative states, respectively (**Fig. 6A**). We identified that APOE4 could affect variability of age-related metabolic change. The FDG PET images generated by average latent features and APOE4 positive and negative status at different ages were subtracted by generated PET of the age of 50 (**Fig. 6B**). We found that metabolic decline in occipital lobe was faster in APOE4 carriers. Distribution of regional metabolism according to APOE4 status was estimated (**Fig. 6C**). Using distribution of metabolic difference between brain metabolism generated by APOE4 status, the significance of difference in regional metabolism was estimated (**fig. S5**). Regional metabolic activity of the calcarine and lingual cortex was significantly higher in APOE4 carrier than APOE4 noncarrier before 60, while that of the hippocampus and amygdala was significantly lower in APOE4 carrier at 50 (**Fig. 6D**). Regional metabolic activity of posterior cingulate, precuneus and caudate, where rapid age-related metabolic decline was found, did not show significant difference in accordance with APOE4 status. Metabolic change in APOE4 carriers and noncarriers of all brain regions was represented with 95% confidence intervals (**fig. S6**).

**Figure 6.**
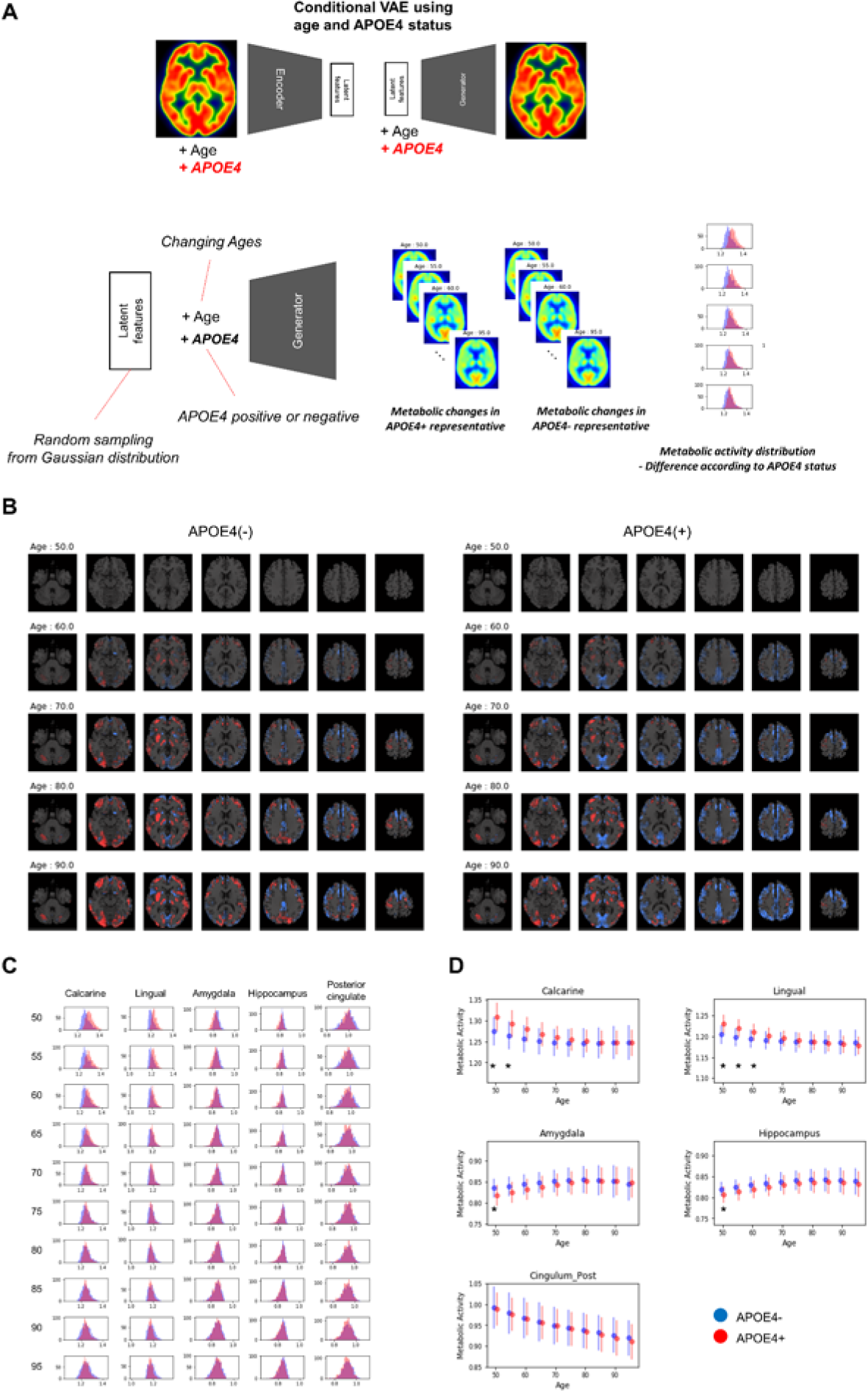
APOE4 status and age-related brain metabolic change. (A) We investigated whether APOE4 status affect age-related metabolic change patterns. A conditional generative model was developed using APOE4 status as well as age. PET images according to different ages were generated for APOE4 carrier and noncarrier, respectively. Resampled features provide distribution difference of brain metabolic topography between APOE4 carriers and noncarreirs. (B) Delta maps were generated by subtracting 50-year-old generated images. Metabolic decline was relatively faster in occipital regions of APOE4 carrier. (C) Histograms of regional metabolic activity were drawn for APOE4 carriers and noncarriers. Before 60, distribution of metabolism of calcarine, lingual cortex, hippocampus and amygdala was different according to APOE4 status. (D) Age-related regional metabolic activity changes were plotted. Red dots represented APOE4 carriers and blue dots represented APOE4 noncarriers. Bars represented standard deviations calculated by the distribution. Nonparametric testing revealed the statistical significance. Asterisks represent uncorrected p < 0.05.

## Discussion

In this study, we predicted aging of brain metabolic topography by using a generative model. Brain metabolic changes are highly variable as aging process and cognitive changes are affected by several individual factors. Our model aimed at generating PET images according to the age trained by cross-sectional PET image data combined with different ages. The model could provide predicted future metabolic decline and validated by real follow-up data. Our results estimate population distribution of normal brain metabolism at each age. This approach was extended to investigate the effect of APOE4 status on the variability of regional brain metabolism at different ages.

Our generative model could find population distribution of brain metabolic topography for each age as well as predict age-related metabolic change. Cognitive aging and age-related functional decrease is accompanied by increased individual variability (*10*). This individual variability is affected by several factors including life experience, genetic backgrounds and susceptibility to neuropathology (*11*). Furthermore, cognitive variability in individuals across time tends to occur mainly after the age of 60 (*12*). Increased individual variability in aging has been supported by several functional neuroimaging studies (*13*–*15*). Nonetheless, age-related brain metabolism change has been briefly estimated by observing overall correlation between age and metabolism (*2*–*6*). This previous approach could not consider individual variability in age-related metabolism (*10*, *16*). Furthermore, it has been difficult to estimate age-dependent normal population distribution of brain image data as the data consist of subjects with different ages. A conventional linear regression model based on overall metabolic changes estimated by all baseline scans failed to estimate personal variability in metabolic decline patterns (**fig. S7**). That was because the model only estimated same decline patterns for all subjects by calculating voxelwise linear regression based on the population.

According to our model, the variability of brain metabolism was represented by the latent features. They determine age-related metabolism patterns because generator used only the latent features and age as inputs. In other words, latent features could reflect individual factors which affect the variability in metabolic topography. Each latent feature represented specific metabolic topography patterns which could be indirectly identified by generating images according to different feature values as shown in **Fig. 5**. In this regard, random resampling of the latent features generated variable brain metabolic topography, which could be used for estimating population distribution. Our result, population distribution of brain metabolism at each age can be applied to quantitatively define regional abnormality in individuals. Using this distribution, we can define how far a given individual brain PET is from the normal population. Thus, this distribution may help to develop quantitative biomarker which represents abnormal aging process of individual brain metabolism.

Our model could predict regional patterns of individual future brain metabolic change, while future prediction of metabolic change was incorrect in quite a few cases. As shown in **fig. S2**, the predicted delta maps were not correlated with real delta maps in individuals at right-lower portions of the matrix. Nonetheless, overall regional metabolic changes obtained by the prediction were highly correlated with those of real follow-up data as shown in **Fig. 2**. That was because VAE eventually extracted age-associated metabolic topography patterns from overall variation of brain metabolism in the training samples. In other words, because of the high variability in age-related brain metabolic changes, VAE-based model generated future brain PET image by approximating global age-related patterns of training samples. It is closely related to the limitation of VAE which tends to generate averaged and blurry images and lack of variety in generated images (*17*). In addition, not only aging but several cognitive, healthy and nutritional factors affect brain metabolic patterns (*18*, *19*). Because of the multiple factors affecting brain metabolism, accurate individual prediction is substantially difficult. In this study, we simply assumed that other factors of future brain PET except age are unchanged. As multiple factors could determine metabolic topography, the generative model with multiple conditions such as cognitive score may improve future PET prediction. Furthermore, combination of another generative model such as generative adversarial model may improve the prediction accuracy (*20*).

Population distribution of metabolic topography revealed that APOE4 carriers showed higher metabolism in the calcarine and lingual cortex, while lower metabolism in the hippocampus and amygdala before 55. The difference in these regions were not found after 60, which suggested that age-related metabolic changes of these regions were greater in APOE4 carriers than noncarriers. The relationship between APOE4 and brain metabolism in normal elderly has been investigated in previous studies as well (*21*). The regions which showed difference metabolism in accordance with APOE4 status were partly different as the previous study showed that metabolic decline was faster in composite region-of-interests including posterior cingulate, precuneus and lateral parietal cortices (*21*). Beside, another study using functional MRI showed APOE4 status affected the differentiation of functional networks including hippocampal and visual networks though they used different modality (*22*). Structural MRI study showed that APOE4 carriers tended to have thicker cortex in temporooccipital areas and steeper age-related decline in cortical thickness (*23*). Although the regions related to APOE4 were partly different according to the studies, our result supports APOE4 carriers could affect functional brain aging patterns. Additionally, by estimating population distribution, we could identify regional metabolic difference at all ages. Our approach can be extended to investigation of the association between other clinical variables and age-related changes. It can eventually help find the factors that determine the individual variability in aging.

To our knowledge, this is the first report that applies a generative model to estimate aging of high dimensional medical data. As an extended application of our approach, PET data according to interpretable features, such as sex and cognitive scores, can be generated by using conditional VAE which aimed at synthesizing virtual data from the conditional distribution (*24*, *25*). This conditional generative model can be used for various problems in neuroimaging analyses. For example, the model may be used for predicting several task-specific functional brain images from a single image data. Virtual task-related brain images can be predicted by inputting tasks as conditional inputs of VAE model. Furthermore, this approach would improve conventional statistical voxelwise analyses of neuroimaging data. An important limitation of the voxelwise analysis is the presence of multiple covariates (*26*, *27*). So far, covariates such as subject’s age and brain volume have been handled as nuisance variables using general linear model. Instead, virtual neuroimaging data in same conditions can be generated by this approach. For instance, we can compare brain images of different groups by generating virtual data with controlled covariates such as same age and brain volume.

As a deep generative model may be able to precisely predict high dimensional data, future application will be extended to various medical implications. Recently, generative models have been used in various biomedical fields as well as neuroimaging data. A generative model was applied to generating novel molecular fingerprints as an artificial intelligence drug discovery framework (*28*). As a recently developed application to medical image processing, generative model was used for automatic lesion segmentation (*29*).

In our study, we predict aging of metabolic topography by generating PET images. In spite of individual variability in age-related change, future regional metabolic changes were precisely predicted. Population distribution of normal brain metabolism at different ages was estimated. It revealed that regional metabolic decline was different according to the APOE4 status. This brain metabolic change prediction method can provide a plausible explanation of individual variability in cognitive aging. Furthermore, we expect that this approach will be extended to the development of preclinical biomarker for several neurodegenerative disorders as well as defining abnormal brain aging.

## Materials and Methods

### Subjects

In this study, the data included subjects recruited in Alzheimer’s Disease Neuroimaging Initiative (ADNI) with FDG PET images (http://adni.loni.usc.edu). The ADNI was launched in 2003 as a public-private partnership, led by Principal Investigator Michael W. Weiner, MD, VA Medical Center and University of California San Francisco. ADNI recruited subjects from over 50 sites across the US and Canada. The primary purpose of ADNI has been to test whether serial imaging and biological markers, and clinical and neuropsychological assessment can be combined to measure the progression of MCI and early AD. For up-to-date information, see http://www.adni-info.org. Written informed consent to cognitive testing and neuroimaging prior to participation was obtained, approved by the institutional review boards of all participating institutions. 393 cognitively normal subjects without Alzheimer’s dementia or mild cognitive impairment performed baseline FDG PET (Age: 73.7 **±** 5.9, range 56.1-90.1). These PET data and their age information were used for developing the model. All subjects underwent clinical and cognitive assessment at the time of acquisition. APOE genotyping was performed on DNA samples obtained from blood. For detailed information on DNA sample preparation and genotyping, see http://www.adni-info.org. For 393 subjects, 113 (28.8%) were APOE4 carriers and 280 (71.2%) were APOE4 noncarriers.

### FDG PET preparation

All the PET images were downloaded from ADNI database. FDG PET images were acquired 30 to 60 min and the images were averaged across the time frames and standardized to have same voxel size (1.5 × 1.5 × 1.5 mm). PET images were acquired in the 57 sites participating in ADNI, scanner-specific smoothing was additionally applied (*30*). PET images were spatially normalized to the Montreal Neurological Institute (MNI) space using statistical parametric mapping (SPM8, www.fil.ion.ucl.ac.uk/spm). Each PET image was divided by mean FDG uptake of the cerebellum for normalization.

### Variational autoencoder for PET volumes

We utilized VAE model to generate virtual PET data according to age information. VAE-based PET image generation is summarized in **Fig. 1A**. VAE is a type of unsupervised learning methods which could represent the high-dimensional data to low-dimensional features. The major strength of the VAE is to generate virtual data from latent features. VAE consisted of two components, encoder and generator. Encoder reduces the dimension of data by compressing them to latent features and generator produces the data from any values of latent features. The generator of VAE is a probabilistic generator which assumes that the data were generated from some conditional distribution and an unobserved variable *z* in latent space. Thus, the probabilistic generator can be defined by *p*_*θ*_(*x*|*z*). θ represents the parameters of generator. The posterior distribution *p*_*θ*_(*z*|*x*) can be obtained by prior distribution *p(z)*,*p*_*θ*_(*z*|*x*)~ *p*(*z*)*p*_*θ*_(*x*|*z*). Variational Bayes learns both parameters, *p*_*θ*_(*x*|*z*) and an approximation *q*_*ϕ*_(*z*|*x*) to the intractable true posterior *p*_*θ*_(*z*|*x*). This is achieved by the loss function,

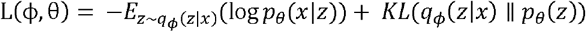

where KL is Kullback-Leibler divergence between the learnt latent distribution and the prior distribution *p*_*θ*_(*z*), acting as a regularization term (*9*). The first term represents reconstruction loss of autoencoder.

In this study, we applied VAE with age information to generate PET image, so used VAE conditioning on another description of the data, *y* (i.e. age information). This model is aimed to generate data from the conditional distribution as well as latent features *z*. Thus, the probabilistic generator and the encoder can be defined by *p*_*θ*_(*x*|*y*,*z*) and *q*_*ϕ*_,(*z*|*x*,*y*), respectively. The loss function is changed to,

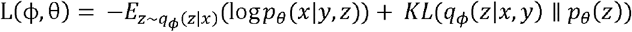

To train VAE, data *X* and age information *y* were encoded into parameters in a latent features *Z*, and decoder network reconstructs data from the latent features and *y* assuming latent features have normal distribution around encoded feature *z*. In practice, generator input was resampled by the encoded latent features *z* assuming normal distribution:*z*_*resampted*_ = *z*_*encoder*_ + *z*_*sd*_ × *ε*, where *ε* represents a random variable (*9*).

### Network architecture and training

To encode 3-dimensional PET volume, we used multiple 3D convolutional layers for encoding. Specific parameters for network architecture are summarized in **fig. S8**. After the multiple convolutional and pooling layers, 3D feature volumes are changed to 1-dimensional features. These features are merged by age information of each subject and additionally connected to hidden layers and, finally, connected to 10 latent features. Accordingly, initial PET volume with 79×95×68 matrix is compressed into 10 dimensional features. Conversely, the generator consists of convolutional and upsampling layers. Upsampling simply repeats each dimension of the data. Input variables of the generator include 10 latent features and age information. The generator decodes these inputs to PET volume.

This conditional VAE model was trained by gradient descent algorithm (Adadelta) (*31*) and took 50 epochs for the training. The VAE was implemented using a deep learning library, Keras (ver. 1.2.2) with Theano (ver. 0.9.0) backend (*32*). 10% of all PET data were used for validation set to determine epoch number and hyperparameters for the neural network architecture.

### Estimation of metabolic activity in brain regions

Regional metabolic activity of brain regions was obtained using predefined volume-of-interests (VOI), automated anatomical labeling (AAL) template. As all PET images were spatially normalized to MNI template, mean metabolic activity value of each brain region was simply obtained by masking specific brain region.

### Prediction of future PET and comparison with follow-up PET

4-year follow-up FDG brain PET scans were obtained in 26 cognitively normal subjects who underwent baseline PET scans. 5-year follow-up FDG brain PET scans were acquired in 11 cognitively normal subjects. Longitudinal change in brain metabolism was evaluated in these subjects. Using baseline PET images of the subjects and age, we generated future PET images. To generate individual future PET image, firstly, baseline PET image was represented into latent features using the encoder. We hypothesized that these latent features were unchanged regardless of subject’s age. 10 latent features of a subject and future age (i.e. baseline age + 4 or 5) were used for generator. We compared real follow-up PET and predicted PET by using delta maps. To measure similarity between predicted and real metabolic changes, voxelwise correlation coefficient was calculated. Similarity measurements were individually obtained. We statistically tested whether other variables including baseline age, gender, APOE4 status, MMSE and follow-up diagnosis affected the prediction of metabolic changes. The similarity measurements, correlation coefficients, of the group according to the APOE4 status, gender and follow-up diagnosis were statistically compared using independent t-test. They were correlated with continuous variables (age and MMSE) using Pearson correlation.

In addition, overall predicted and real regional changes were calculated by AAL map. Overall regional metabolic change was calculated by mean value across all subjects. The correlation between regional metabolic changes of predicted and real follow-up PET across brain regions was tested by Pearson correlation. For visualizing the similarity between predicted and real metabolic changes, Bland-Altman plots were drawn. 95% confidence interval for error of predicted regional metabolic change was calculated.

### Generation of age-related metabolic change movie

The overall age-related metabolic change pattern was evaluated by the generator model. Firstly, PET data of all subjects were represented by 10 latent features using encoder. The mean feature values were entered into the generator with different age information between age of 50 and 100. Thus, we could obtain representative PET image of each age. To visualize age-related metabolic change, we generate subtraction map. Generated PET images with different age was subtracted by a representative brain PET generated by age of 50. These subtraction maps were also visualized by an animation.

### Population distribution of regional metabolic activity at each age

We estimated population distribution of regional metabolic activity by resampling generated PET images. 10 latent features were randomly resampled assuming each latent feature has normal distribution. Mean and standard deviation of each latent feature were determined by the feature values of all subjects. 1000 resampled brain PET images were generated and regional metabolic activity was obtained. Population distribution of metabolic activity of each region was drawn by histograms and age-related changes with confidence intervals were drawn.

### Metabolic topography according to latent features

To assess the relationship between latent features and brain metabolic patterns, brain PET images were generated by changing values of the latent features. Mean values of latent features were used for generating PET except two features for estimating effects on brain metabolism. These two features were changed from -2.0 to 2.0 and generated virtual PET images for plotting.

### Variability in age-related metabolic change according to APOE4 status

To evaluate age-related metabolic change patterns according to the APOE4 status, another VAE model was trained. Conditional VAE with age and APOE4 status information was used, so, conditional variable, *y*, includes age and APOE4 status as different dimensions. The training process and network architectures were same with conditional VAE with age only.

The overall age-related metabolic change patterns according to APOE4 status was evaluated as population distribution estimation. Randomly resampled latent features and different age values were entered into the generator with each APOE4 status respectively. PET images of each age and APOE4 status were generated and regional metabolic activity was obtained by predefined regions. Population distribution of regional metabolic activity was estimated for APOE4 carriers and noncarriers. To find statistically different regions, we calculated the difference between regional metabolic activity generated by APOE4 carriers and noncarriers. To define statistical significance, p-values were computed by distribution of the difference. They were proportion values that represented the difference was less than or more than 0. Brain regions with different metabolic activity were found at each age. The difference with uncorrected p-value less than 0.05 was regarded as significant brain regions.

## Supplementary Materials

Fig. S1. Voxelwise correlation between predicted and real metabolic changes.

Fig. S2. Similarity between predicted and real metabolic changes for individual subjects.

Fig. S3. Overall brain metabolism aging patterns.

Fig. S4. Regional metabolic changes of all brain regions.

Fig. S5. Distribution of difference of regional metabolic activity between APOE4 carriers and noncarriers.

Fig. S6. Age-related metabolic changes according to APOE4 status.

Fig. S7. Age-related metabolic change estimated by linear regression.

Fig. S8. Network architecture of variational autoencoder model.

## Acknowledgement

In this study, the data included subjects recruited in Alzheimer’s Disease Neuroimaging Initiative (ADNI) with FDG PET images (http://adni.loni.usc.edu)

## REFERENCES

1. W. J. Jagust, S. Landau, L. Shaw, J. Trojanowski, R. Koeppe, E. Reiman, N. Foster, R. C. Petersen, M. Weiner, J. Price, Relationships between biomarkers in aging and dementia. Neurology 73, 1193–1199 (2009).

2. R. Duara, C. Grady, J. Haxby, D. Ingvar, L. Sokoloff, R. A. Margolin, R. G. Manning, N. R. Cutler, S. I. Rapoport, Human brain glucose utilization and cognitive function in relation to age. Ann Neurol 16, 703–713 (1984).

3. J. R. Moeller, T. Ishikawa, V. Dhawan, P. Spetsieris, F. Mandel, G. E. Alexander, C. Grady, P. Pietrini, D. Eidelberg, The metabolic topography of normal aging. J Cereb Blood Flow Metab 16, 385–398 (1996).

4. M. C. Petit-Taboue, B. Landeau, J. F. Desson, B. Desgranges, J. C. Baron, Effects of healthy aging on the regional cerebral metabolic rate of glucose assessed with statistical parametric mapping. Neuroimage 7, 176–184 (1998).

5. A. Loessner, A. Alavi, K. U. Lewandrowski, D. Mozley, E. Souder, R. E. Gur, Regional cerebral function determined by FDG-PET in healthy volunteers: normal patterns and changes with age. J Nucl Med 36, 1141–1149 (1995).

6. D. Yanase, I. Matsunari, K. Yajima, W. Chen, A. Fujikawa, S. Nishimura, H. Matsuda, M. Yamada, Brain FDG PET study of normal aging in Japanese: effect of atrophy correction. Eur J Nucl Med Mol Imaging 32, 794–805 (2005).

7. N. Raz, U. Lindenberger, K. M. Rodrigue, K. M. Kennedy, D. Head, A. Williamson, C. Dahle, D. Gerstorf, J. D. Acker, Regional brain changes in aging healthy adults: general trends, individual differences and modifiers. Cerebral cortex 15, 1676–1689 (2005).

8. C. Grady, The cognitive neuroscience of ageing. Nature reviews. Neuroscience 13, 491–505 (2012).

9. D. P. Kingma, M. Welling, Auto-encoding variational bayes. arXiv preprint arXiv:1312.6114, (2013).

10. R. Ylikoski, A. Ylikoski, P. Keskivaara, R. Tilvis, R. Sulkava, T. Erkinjuntti, Heterogeneity of cognitive profiles in aging: successful aging, normal aging, and individuals at risk for cognitive decline. Eur J Neurol 6, 645–652 (1999).

11. P. Shammi, E. Bosman, D. T. Stuss, Aging and variability in performance. Aging, Neuropsychology, and Cognition 5, 1–13 (1998).

12. R. S. Wilson, L. A. Beckett, L. L. Barnes, J. A. Schneider, J. Bach, D. A. Evans, D. A. Bennett, Individual differences in rates of change in cognitive abilities of older persons. Psychol Aging 17, 179–193 (2002).

13. E. L. Glisky, S. R. Rubin, P. S. Davidson, Source memory in older adults: an encoding or retrieval problem? J Exp Psychol Learn Mem Cogn 27, 1131–1146 (2001).

14. M. D’Esposito, L. Y Deouell, A. Gazzaley, Alterations in the BOLD fMRI signal with ageing and disease: a challenge for neuroimaging. Nature Reviews Neuroscience 4, 863–872 (2003).

15. A. Z. Burzynska, C. N. Wong, M. W. Voss, G. E. Cooke, E. McAuley, A. F. Kramer, White matter integrity supports BOLD signal variability and cognitive performance in the aging human brain. PLoS One 10, e0120315 (2015).

16. D. S. Knopman, C. R. Jack,Jr., H. J. Wiste, E. S. Lundt, S. D. Weigand, P. Vemuri, V. J. Lowe, K. Kantarci, J. L. Gunter, M. L. Senjem, M. M. Mielke, R. O. Roberts, B. F. Boeve, R. C. Petersen, 18F-fluorodeoxyglucose positron emission tomography, aging, and apolipoprotein E genotype in cognitively normal persons. Neurobiol Aging 35, 2096–2106 (2014).

17. A. Dosovitskiy, T. Brox, in Advances in Neural Information Processing Systems. (2016), pp. 658–666.

18. M. Belanger, I. Allaman, P. J. Magistretti, Brain energy metabolism: focus on astrocyte-neuron metabolic cooperation. Cell Metab 14, 724–738 (2011).

19. S. Cunnane, S. Nugent, M. Roy, A. Courchesne-Loyer, E. Croteau, S. Tremblay, A. Castellano, F. Pifferi, C. Bocti, N. Paquet, H. Begdouri, M. Bentourkia, E. Turcotte, M. Allard, P. Barberger-Gateau, T. Fulop, S. I. Rapoport, Brain fuel metabolism, aging, and Alzheimer’s disease. Nutrition 27, 3–20 (2011).

20. I. Goodfellow, J. Pouget-Abadie, M. Mirza, B. Xu, D. Warde-Farley, S. Ozair, A. Courville, Y Bengio, in Advances in neural information processing systems. (2014), pp. 2672–2680.

21. H. Oh, C. Habeck, C. Madison, W. Jagust, Covarying alterations in Abeta deposition, glucose metabolism, and gray matter volume in cognitively normal elderly. Hum Brain Mapp 35, 297–308 (2014).

22. A. J. Trachtenberg, N. Filippini, K. P. Ebmeier, S. M. Smith, F. Karpe, C. E. Mackay, The effects of APOE on the functional architecture of the resting brain. Neuroimage 59, 565–572 (2012).

23. T. Espeseth, L. T. Westlye, A. M. Fjell, K. B. Walhovd, H. Rootwelt, I. Reinvang, Accelerated age-related cortical thinning in healthy carriers of apolipoprotein E epsilon 4. Neurobiol Aging 29, 329–340 (2008).

24. K. Sohn, H. Lee, X. Yan, in Advances in Neural Information Processing Systems. (2015), pp. 3483–3491.

25. D. P. Kingma, S. Mohamed, D. J. Rezende, M. Welling, in Advances in Neural Information Processing Systems. (2014), pp. 3581–3589.

26. K. J. Friston, A. P. Holmes, K. J. Worsley, J. P. Poline, C. D. Frith, R. S. Frackowiak, Statistical parametric maps in functional imaging: a general linear approach. Human brain mapping 2, 189–210 (1994).

27. K. M. Petersson, T. E. Nichols, J.-B. Poline, A. P. Holmes, Statistical limitations in functional neuroimaging. I. Non-inferential methods and statistical models. Philosophical Transactions of the Royal Society of London B: Biological Sciences 354, 1239–1260 (1999).

28. A. Kadurin, A. Aliper, A. Kazennov, P. Mamoshina, Q. Vanhaelen, K. Khrabrov, A. Zhavoronkov, The cornucopia of meaningful leads: Applying deep adversarial autoencoders for new molecule development in oncology. Oncotarget, (2016).

29. V Alex, M. S. Kp, S. S. Chennamsetty, G. Krishnamurthi, in SPIE Medical Imaging. (International Society for Optics and Photonics, 2017), pp. 101330G-101330G–101339.

30. W. J. Jagust, S. M. Landau, R. A. Koeppe, E. M. Reiman, K. Chen, C. A. Mathis, J. C. Price, N. L. Foster, A. Y. Wang, The Alzheimer’s Disease Neuroimaging Initiative 2 PET Core: 2015. Alzheimers Dement 11, 757–771 (2015).

31. M. D. Zeiler, ADADELTA: an adaptive learning rate method. arXiv preprint arXiv:1212.5701, (2012).

32. F. Bastien, P. Lamblin, R. Pascanu, J. Bergstra, I. Goodfellow, A. Bergeron, N. Bouchard, D. Warde-Farley, Y. Bengio, Theano: new features and speed improvements. arXiv preprint arXiv:1211.5590, (2012).

